# The evolution of skin pigmentation associated variation in West Eurasia

**DOI:** 10.1101/2020.05.08.085274

**Authors:** Dan Ju, Iain Mathieson

## Abstract

Skin pigmentation is a classic example of a polygenic trait that has experienced directional selection in humans. Genome-wide association studies have identified well over a hundred pigmentation-associated loci, and genomic scans in present-day and ancient populations have identified selective sweeps for a small number of light pigmentation-associated alleles in Europeans. It is unclear whether selection has operated on all the genetic variation associated with skin pigmentation as opposed to just a small number of large-effect variants. Here, we address this question using ancient DNA from 1158 individuals from West Eurasia covering a period of 40,000 years combined with genome-wide association summary statistics from the UK Biobank. We find a robust signal of directional selection in ancient West Eurasians on skin pigmentation variants ascertained in the UK Biobank, but find this signal is driven mostly by a limited number of large-effect variants. Consistent with this observation, we find that a polygenic selection test in present-day populations fails to detect selection with the full set of variants; rather, only the top five show strong evidence of selection. Our data allow us to disentangle the effects of admixture and selection. Most notably, a large-effect variant at SLC24A5 was introduced to Europe by migrations of Neolithic farming populations but continued to be under selection post-admixture. This study shows that the response to selection for light skin pigmentation in West Eurasia was driven by a relatively small proportion of the variants that are associated with present-day phenotypic variation.

**Significance:** Some of the genes responsible for the evolution of light skin pigmentation in Europeans show signals of positive selection in present-day populations. Recently, genome-wide association studies have highlighted the highly polygenic nature of skin pigmentation. It is unclear whether selection has operated on all of these genetic variants or just a subset. By studying variation in over a thousand ancient genomes from West Eurasia covering 40,000 years we are able to study both the aggregate behavior of pigmentation-associated variants and the evolutionary history of individual variants. We find that the evolution of light skin pigmentation in Europeans was driven by frequency changes in a relatively small fraction of the genetic variants that are associated with variation in the trait today.

## Introduction

Skin pigmentation exhibits a gradient of variation across human populations that tracks with latitude (1). This gradient is thought to reflect selection for lighter skin pigmentation at higher latitudes, as lower UVB exposure reduces vitamin D biosynthesis which affects calcium homeostasis and immunity (2–4). Studies of present-day and ancient populations have revealed signatures of selection at skin pigmentation loci (5–9), and single nucleotide polymorphisms (SNPs) associated with light skin pigmentation at some of these genes exhibit a signal of polygenic selection in Western Eurasians (10). However, this observation, the only documented signal of polygenic selection for skin pigmentation, is based on just four loci (*SLC24A5*, *SLC45A2*, *TYR*, and *APBA2/OCA2*).

Therefore, while the existence of selective sweeps at a handful of skin pigmentation loci is well-established, the evidence for polygenic selection–a coordinated shift in allele frequencies across many trait-associated variants (11)–is less clear. Recently, genome-wide association studies (GWAS) of larger samples and more diverse populations (12–15) have emphasized the polygenic architecture of skin pigmentation. This raises the question of whether selection on skin pigmentation acted on all of this variation or rather was driven by selective sweeps at a relatively small number of loci.

The impact of demographic transitions on the evolution of skin pigmentation also remains an open question. The Holocene history (∼12,000 years before present (BP)) of Europe was marked by waves of migration and admixture between three highly diverged populations: hunter-gatherers, Early Farmers, and Steppe-ancestry populations (16, 17). Skin pigmentation-associated loci may have been selected independently or in parallel in one or more of these source populations or may instead only have been selected after admixture. The impact of ancient shifts in ancestry is difficult to resolve using present-day data, but using ancient DNA, we can separate the effects of ancestry and selection and identify which loci were selected in which populations.

Here we use ancient DNA to track the evolution of loci that are associated with skin pigmentation in present-day Europeans. Although we cannot make predictions about the phenotypes of ancient peoples, we can assess the extent to which they carried the same light pigmentation alleles that are present today. This allows us to identify which pigmentation-associated variants have changed in frequency due to positive selection and the timing of these selective events. We present the first systematic survey of the evolution of European skin pigmentation-associated variation, tracking over a hundred loci over 40,000 years of human history—almost the entire range of modern human occupation of Europe.

## Results

### Skin pigmentation associated SNPs show signal of positive selection in ancient West Eurasians

We obtained skin pigmentation-associated SNPs from the UK Biobank GWAS for skin colour released by the Neale Lab (14). We analyzed the evolution of these variants using two datasets of publicly available ancient DNA (Fig. 1 *A-C*, Fig. S1). The first (“capture-shotgun”) consists of data from 1158 individuals dating from 45,000 to 715 years BP genotyped at ∼1.2 million variants (7, 16, 26–35, 18, 36–45, 19, 46–52, 20–25). The second (“shotgun”) is a subset of 248 individuals with genome-wide shotgun sequence data (16, 18, 31–33, 39, 42, 45–49, 20, 50, 53–60, 22, 24, 26–30). This smaller dataset allows us to capture variants that may not be well-tagged by the genotyped SNPs in the capture-shotgun dataset. Weighting variants by their GWAS-estimated effect sizes, we show that the polygenic score–the weighted proportion of dark pigmentation alleles–decreased significantly over the past 40,000 years (Fig. 1 *A* & *B*, *P* < 1×10^−4^ based on a genomic null distribution and accounting for changes in ancestry; Fig. S2). The genetic scores of the oldest individuals in the dataset fall within the range of present-day West African populations, showing that Early Upper Paleolithic [∼50-20,000 years BP] modern humans, such as Ust’Ishim, carried few of the light skin pigmentation alleles that are common in present-day Europe.

**Figure 1.**
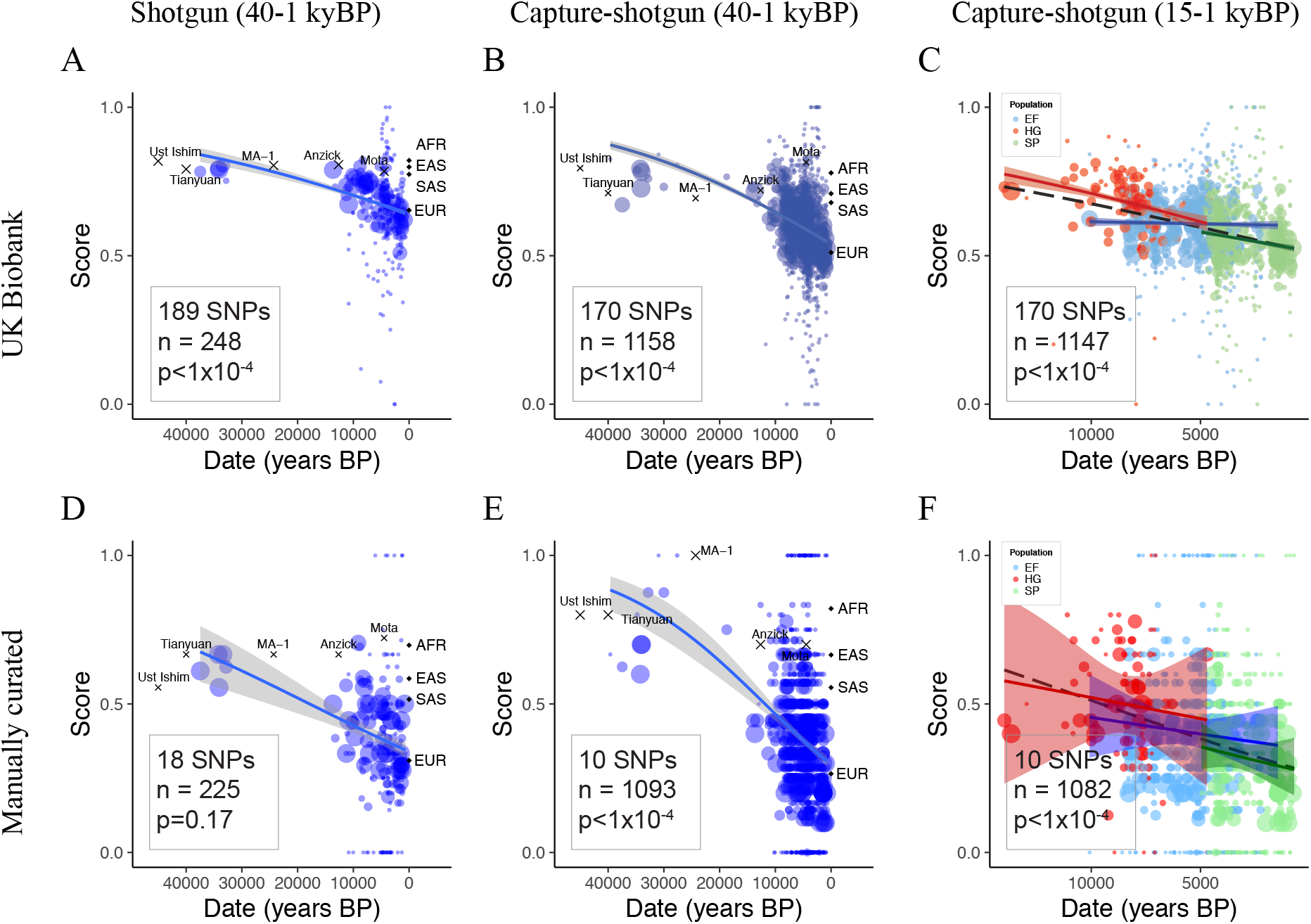
Genetic scores for skin pigmentation over time. Solid lines indicate fitted mean scores with gray 95% confidence intervals. Weighted scores based on UK Biobank SNPs and unweighted scores based on manually curated SNPs for samples in the (A,D) shotgun dataset, (B,E) capture-shotgun dataset, and (C,F) capture-shotgun dataset dated within the past 15,000 years. Scores of labelled samples are plotted but not included in regressions. Averages for 1000 Genomes super-populations are plotted at time 0. Area of points scale with number of SNPs genotyped in the individual.

Over the past 15,000 years, which covers most of the data, the polygenic score decreased (*P* < 1×10^−4^) at a similar rate to the estimated rate over the 40,000 year period (5.27 × 10^−5^ per-year vs 3.47 × 10^−5^ per-year; *P* = 0.04). Stratifying individuals by ancestry (hunter-gatherer, Early Farmer, and Steppe) using ADMIXTURE revealed baseline differences in genetic score across groups (Fig. 1*C*). Mesolithic hunter-gatherers carried fewer light pigmentation alleles than Early Farmers or Steppe ancestry populations. This difference (0.091 score units) was similar to the difference between Mesolithic hunter-gatherers and Early Upper Paleolithic populations (0.097 score units). We tested for evidence of change in score over time within groups, finding no significant change within hunter-gatherer or Early Farmer populations (*P* = 0.24; *P* = 0.29). Steppe ancestry populations, however, exhibited a significant decrease, even after accounting for changes in ancestry over time (*P* = 0.008), with a steeper slope than that of Early Farmer or hunter-gatherer (*P* = 0.004 and *P* = 0.08, respectively).

While the UK Biobank is well-powered to detect variation that is segregating in present-day Europeans, it may not detect variants that are no longer segregating. We therefore manually curated a separate list of 18 SNPs identified as associated with skin pigmentation from seven studies of populations representing diverse ancestries (Table S6). Using this set of SNPs to generate unweighted scores, we observed similar results as with the UK Biobank ascertained SNPs (Fig. 1 *D-F*). We find a significant decrease in genetic score over both 40,000 and 15,000 years in the capture-shotgun dataset (*P* < 1 × 10^−4^ for both), although the decrease in the shotgun dataset was not significant (*P* = 0.17), probably reflecting a lack of power from the small number of ancient samples and SNPs.

### Ancestry and selection have different effects on the histories of each variant

Next, we investigated the evolution of each variant independently. We separated the effects of ancestry and selection in the data by regressing presence or absence of the alternate allele for each pigmentation-associated SNP on both date and ancestry, inferred using PCA (61). Significant effects of the ancestry on frequency can be interpreted as changes in allele frequency that can be explained by changes in ancestry. Significant changes in frequency over time, after accounting for ancestry, can be interpreted as evidence for selection in West Eurasia occurring during the analyzed time period.

The SNP rs16891982 at the *SLC45A2* locus shows the strongest evidence for selection across analyses (Fig. 2), but we note this does not necessarily imply positive selection over the entire time transect. We observed little evidence for demographic transitions in driving the increase in light allele frequency over time for this SNP. In addition, we found robust signals of selection for the light allele of rs1126809 near *TYR*, rs12913832 and rs1635168 near *HERC2*, and rs7109255 near *GRM5* (Fig. 2 *B* & *C*). In some cases–for example at rs6120849 near *EDEM2* and rs7870409 near *RALGPS1*–light alleles decreased in frequency more than can be explained by changes in ancestry (Fig. 2 *B* & *C*). The dark allele at rs6120849 (*EDEM2*) is also associated with increased sitting and standing height in the UK Biobank, and the dark allele of rs7870409 (*RALGPS1*) is associated with increased arm and trunk fat percentage (62). These observations raise the possibility of pleiotropic effects constraining the evolution of these loci or of spurious pleiotropy due to uncorrected population structure in the GWAS.

**Figure 2.**
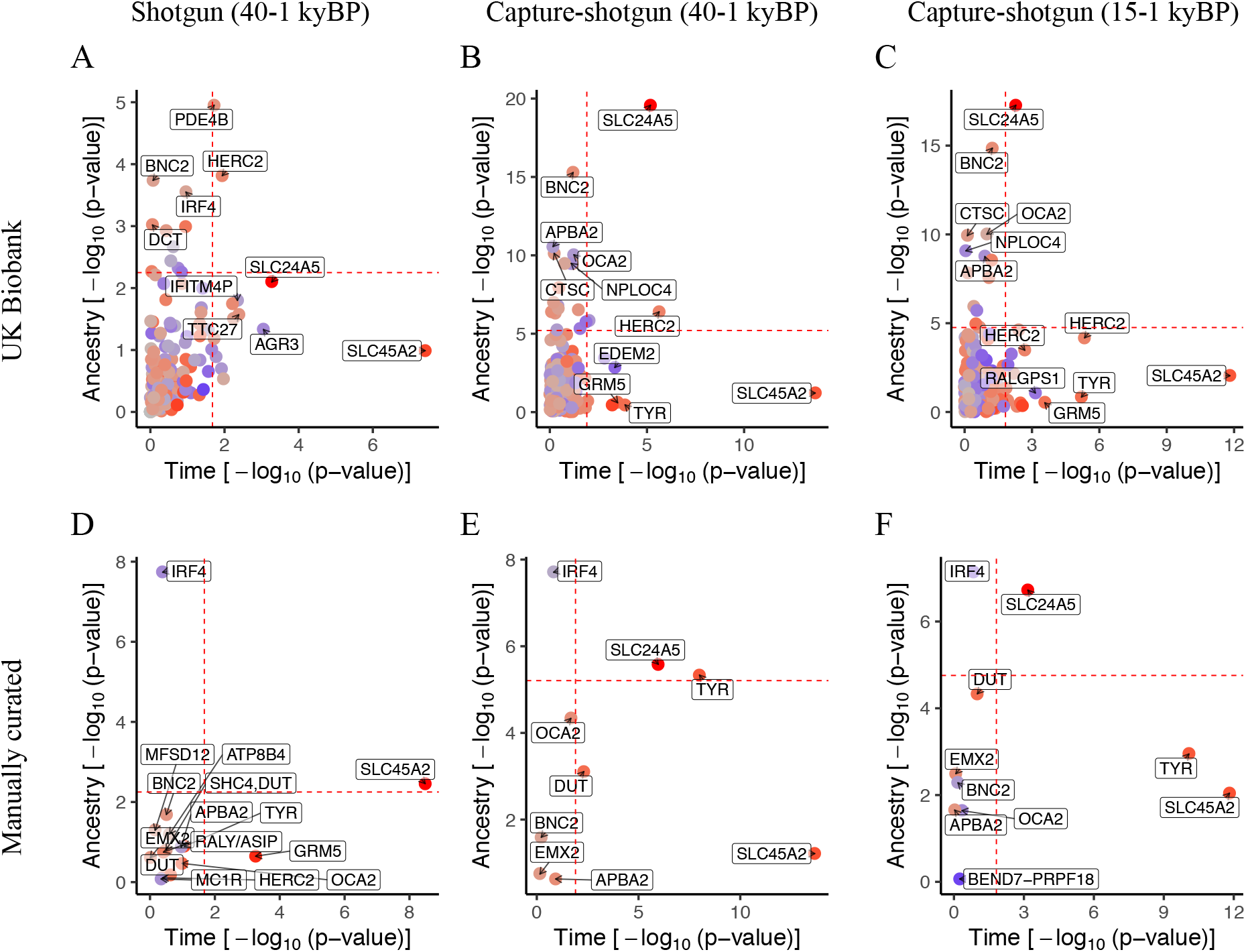
Plot of *P*-values for ancestry and time for individual skin pigmentation SNPs. Dashed lines indicate fifth percentile of *P*-values. UK Biobank SNPs using the shotgun dataset are plotted in (A) and capture-shotgun dataset in (B,C). Manually curated SNPs using the shotgun dataset are plotted in (D) and capture-shotgun dataset in (E,F). SNPs are colored according to whether the light allele is increasing over time (red) or decreasing (blue), with saturation determined by the magnitude of change.

On the other hand, the changes in frequencies of several variants appear to have been driven largely by changes in ancestry. For example, rs4778123 near *OCA2*, rs2153271 near *BNC2* and rs3758833 near *CTSC* show evidence that their changes in frequency were consistent with changes in genome-wide ancestry (Fig. 2 *B* & *C*). At *SLC24A5*, rs2675345 shows evidence of change with both ancestry and time, suggesting that even after the spread of the light allele from migrating populations in the Neolithic (7), selection continued to occur post-admixture. Again, not all of these cases involve an increase in the frequency of the light pigmentation allele over time. The light allele of rs4778123 (*OCA2*) was at high frequency in hunter-gatherers but lower in later populations (Fig. S4). From the manually curated set of SNPs (Fig. 2 *D-F*), rs12203592 near *IRF4* also displays a marked effect of ancestry with higher light allele frequency in hunter-gatherers as well. While rs12203592 was not present in the UK Biobank summary statistics, another SNP at the *IRF4* locus, rs3778607, was present but with a lower ancestry effect (Fig S5).

### Lack of PBS selection signal across pigmentation SNPs in source populations of Europeans

As we described in the previous section, the frequencies of several skin pigmentation SNPs appear to have been driven by changes in ancestry. This observation suggests that their frequencies have diverged between the European source populations and then subsequently been driven by admixture. Frequency differences across the source populations might reflect the action of either genetic drift or selection. To examine this question, we looked for evidence of selection at skin pigmentation SNPs in each ancestral population using the population branch statistic (PBS) test (63). We assigned ancient individuals to the groups hunter-gatherer, Early Farmer, and Steppe ancestry based on unsupervised ADMIXTURE with K=3 (Fig. S6) and used the observed frequencies in each group to calculate PBS in non-overlapping 50-SNP windows.

Few of the skin pigmentation loci show extreme PBS values (Fig. 3, Fig. S7). On the Early Farmer and Steppe ancestry branches, but not hunter-gatherer branch, the *SLC24A5* locus exhibited the strongest signal (Fig. 3 *B* & *C*), indicating selection at the locus in both Early Farmer and Steppe source populations. The *RABGAP1* locus also exhibited an elevated signal of selection in Early Farmer and Steppe, which remained when using a 20-SNP window size (Fig. S8 *B* and *C*). However, unlike the *SLC24A5* variant, this SNP (rs644490) did not show a substantial effect of ancestry from the individual SNP regression analysis. Across the ancestral groups, we observed an elevated PBS at the *OCA2* locus around rs9920172, with the most extreme value on the hunter-gatherer branch. As expected, this SNP does not show an effect of ancestry in allele frequency change over time. As a whole, the PBS values of the 170 skin pigmentation SNPs do not significantly deviate from the genome-wide distribution of PBS for any of the ancient source populations (Fig. S9).

**Figure 3.**
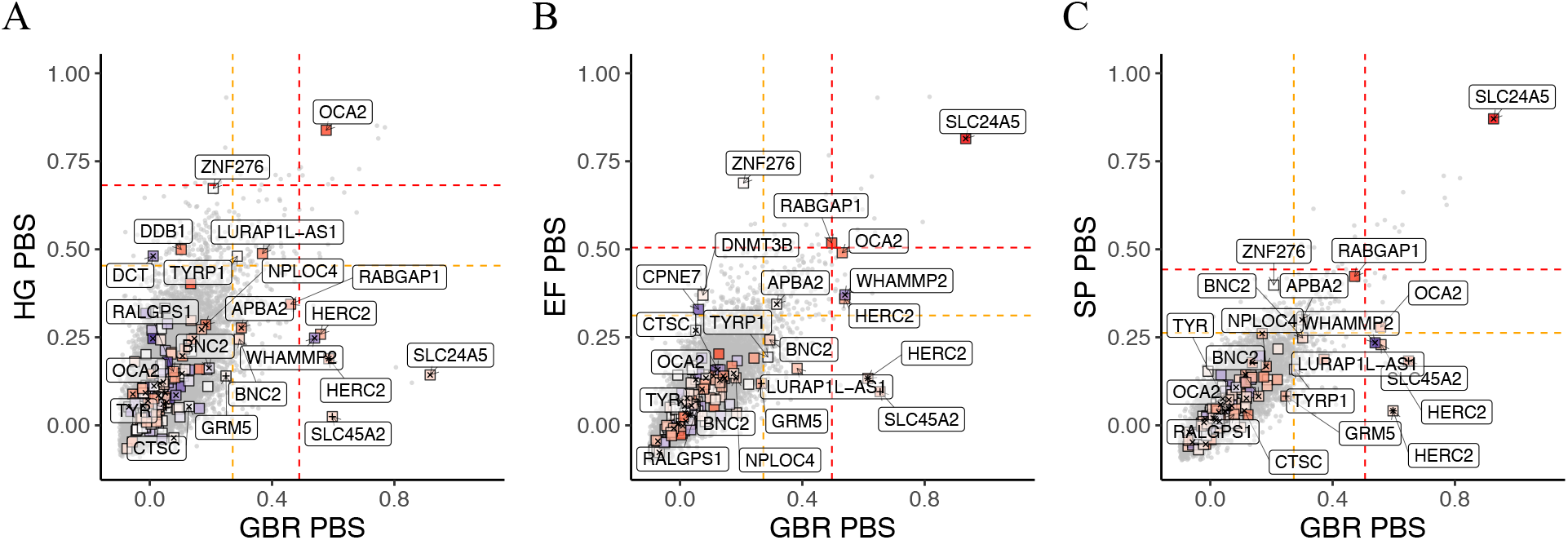
Joint PBS distributions for 40-SNP windows across the genome for GBR-CHB-YRI on the x-axis and X-CHB-YRI on the y-axis, with X being (A) hunter-gatherer, (B) Early Farmer, and (C) Steppe. Boxes represent windows centered around UK Biobank skin pigmentation SNPs and are colored redder according to how much higher the light allele frequency is in X compared to YRI. Plusses (+) mark windows around SNPs at the extremes of the time *P*-value distribution from Fig. 2, whereas crosses (X) mark windows around SNPs at the extremes of the ancestry distribution. Nearest genes are labelled. Orange and red lines represent top 1 and 0.1 percentiles.

### Signals of selection are restricted to the SNPs with the largest effect sizes

Since our results suggest that pigmentation-associated variants exhibit different changes in allele frequency, we further examined the signal of selection found in Fig. 1. In general, SNPs with larger GWAS effect sizes changed more in frequency over time than those with small effect sizes (*P* = 2.8 × 10^−18^; adjusted R^2^=0.36) (Fig. S10*A*). To understand the contribution of these large GWAS effect size SNPs, we iteratively removed SNPs from the polygenic score calculations (Fig 4*A*). Removing the SNP rs16891982 at *SLC45A2*, we still found a significant decrease of score over time (*P* < 1 × 10^−4^), but the estimated rate of change (*β*_*time*_) decreased by 39 percent. Removing the top two SNPs at the *SLC45A2* and *SLC24A5* locus further attenuated the signal with a decrease in *β*_*time*_ of 58 percent (*P* = 3.96 × 10^−4^). We removed the five SNPs with the largest GWAS effect sizes and found the effect of time attenuated (*P* = 0.026) with a decrease in *β*_*time*_ of 78 percent. Removing the top 10 SNPs attenuated the signal even more (*P* = 0.050) with an 81 percent decrease and removal of 15 or 20 SNPs essentially abolished the signal (*P* = 0.147 and *P* = 0.52).

**Figure 4.**
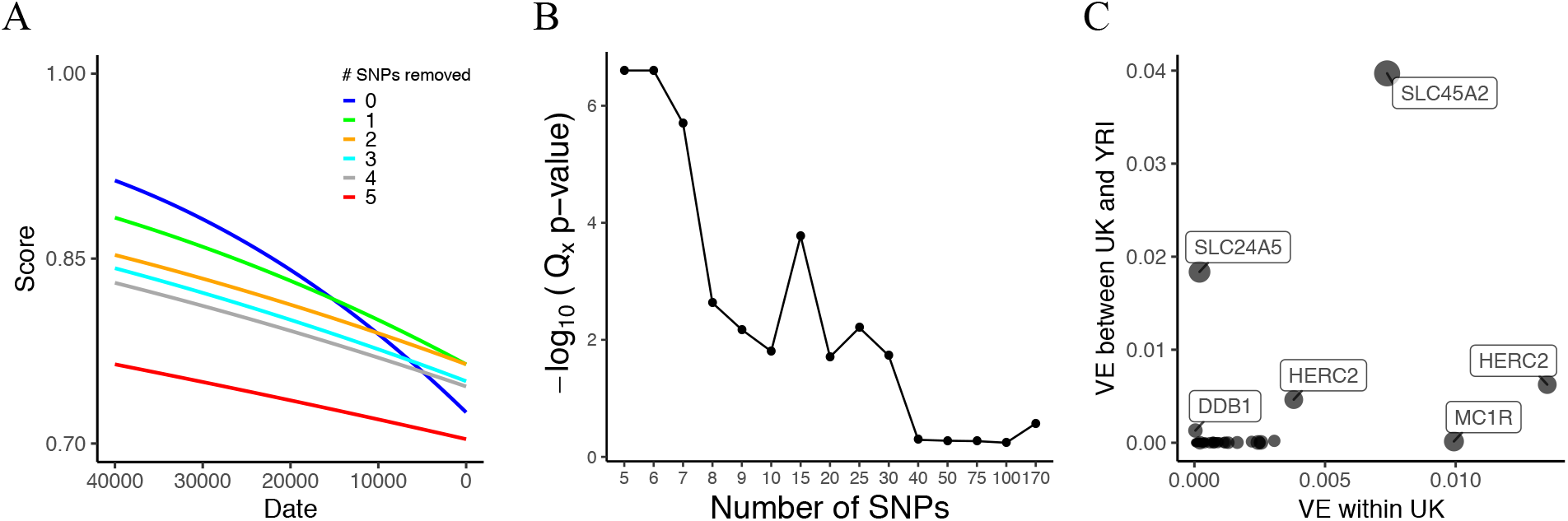
(A) Regressions of genetic score based on UK Biobank SNPs using the capture-shotgun dataset over date, with scores using all SNPs and iteratively removing top GWAS effect size SNPs. (B) *Q_x_* empirical −log_10_(*P*-values) using UK Biobank skin pigmentation associated variants with all 1000 Genomes populations. Different numbers of SNPs were used to calculate *Q_x_* which were ordered by GWAS-estimated effect size. (C) Variance explained by UK Biobank SNPs with nearest gene labelled between UK and YRI populations (y-axis) and within the UK population (x-axis).

Using present-day populations from the 1000 Genomes Project (64), we also tested for polygenic selection with the *Q_x_* statistic (10), which measures overdispersion of trait-associated SNPs relative to the genome-wide expectation. We performed the test with different numbers of SNPs ordered by largest to smallest absolute GWAS effect size (Fig. 4*B*). Using all 170 SNPs in the test, we failed to observe a signal of polygenic selection (*P* = 0.27). However, when we restricted to the top 5 largest GWAS effect size SNPs, we found a significant signal (*P* < 2.5 × 10^−7^). We observed a significant Q_x_ statistics at *P* < 0.05 up to the top 30 SNPs, but it appears this signal is driven by the top SNPs of the largest effect. After removing the top 5 SNPs and testing the remaining 25 top SNPs, the signal of polygenic selection disappeared (*P* = 0.59). Examining the predicted variance explained per SNP (Fig. 4*C*, Fig. S11) we predict that of the UK Biobank SNPs two SNPs at *SLC45A2* and *SLC24A5* contribute substantially to the difference between West African and European populations. Most UK Biobank skin pigmentation-associated SNPs thus are predicted to contribute relatively little to the between-population variance.

## Discussion

Large whole-genome ancient DNA datasets have, in the past few years, allowed us to track the evolution of variation associated with both simple and complex traits (65, 66). The majority of samples are from Western Eurasia and we focus on that region, noting that a parallel process of selection for light skin pigmentation has acted in East Asian populations (9, 67). In West Eurasia, selection for skin pigmentation appears to have been dominated by a small number of selective sweeps at large-effect variants. There are a number of possible explanations for this. Many of the variants detected by the UK Biobank GWAS may have such small effects that they were effectively neutral or experienced such weak selection that we cannot detect it. Other variants may not have been selected because of pleiotropic constraint. Finally, GWAS effect size estimates in European populations today may not reflect the impact these variants had in the past due to epistatic or gene-environment interactions. This would not affect the individual SNP time series and PBS analyses since they do not rely on effect sizes as weights but would reduce power for the weighted polygenic score and *Q_x_* analyses.

We find little evidence of parallel selection on independent haplotypes at skin pigmentation loci, suggesting that that differences in allele frequency across ancestry groups were mostly due to genetic drift. One exception is that the light allele at *SLC24A5* was nearly fixed in both Early Farmer and Steppe ancestry populations due to selection. However, even for this variant we observe a signal of ongoing selection in our data even after admixture with hunter-gatherers, indicating continued selection after admixture. This is analogous to the rapid selection at the same locus for the light allele introduced via admixture into the KhoeSan, who now occupy southern Africa (12, 68).

We are also able to test previous claims about selection on particular pigmentation genes. In contrast to reports of positive selection in Europeans at the *MC1R* locus (69, 70), we find no evidence of positive selection in Europeans. Among UK Biobank SNPs, rs1805007 near *MC1R* explains a relatively large amount of variation within the UK but is predicted to explain relatively little of the variation between Europe and West Africa (Fig. 4*C*). The TYRP1 locus has been previously identified as a target of selection in Europeans (5, 8, 9, 71, 72) although some studies (73, 74) have questioned this finding. Our analysis supports the hypothesis of selection at this locus, with a window centered around rs1325132 in the top 0.77% of genome-wide PBS windows on the hunter-gatherer lineage. At the *KITLG* locus, we find some support for a shared selective sweep in East Asians and Europeans (9, 75, 76). Some studies (73, 77) detect a signal of more recent selection in Europeans at the *KITLG* SNP rs12821256 that is functionally associated with blond hair color (78). We find no evidence of selection on the West Eurasian lineage at this SNP, but this may reflect a lack power to detect selection on a relatively rare variant. Finally, at the *OCA2* locus, we recapitulate an observation of independent selection in Europeans and East Asians (67, 79) (Fig. S7). In the ancient populations, we found an elevated PBS signal around rs9920172 that was most prominent in hunter-gatherers but also elevated in Early Farmer and Steppe ancestry populations (Fig. 3), suggesting a relatively early episode of selection in West Eurasians.

Relatively dark skin pigmentation in Early Upper Paleolithic Europe would be consistent with those populations being relatively poorly adapted to high latitude conditions as a result of having recently migrated from lower latitudes. On the other hand, although we have shown that these populations carried few of the light pigmentation alleles that are segregating in present-day Europe, they may have carried different alleles that we cannot now detect. As an extreme example, Neanderthals and the Altai Denisovan individual show genetic scores that are in a similar range to Early Upper Paleolithic individuals (Table S1), but it is highly plausible that these populations, who lived at high latitudes for hundreds of thousands of years, would have adapted independently to low UV levels. For this reason, we cannot confidently make statements about the skin pigmentation of ancient populations.

Our study focused, for reasons of data availability, on the history of skin pigmentation evolution in West Eurasia. But there is strong evidence that a parallel trend of adaptation to low UVB conditions occurred in East Asia (67, 80–82). Less is known about the loci that have been under selection in East Asia, aside from some variants at *OCA2* (79, 80, 82). Similarly, multiple studies have documented selection for both lighter and darker skin pigmentation in parts of Africa (12, 13, 83). Future work should test whether the process of adaptation in other parts of the world was similar to that in Europe. The lack of known skin pigmentation loci in scans of positive selection in East Asian populations (84, 85) raises the possibility that selection may have been more polygenic in East Asians than in Europeans. Finally, the evidence for polygenic directional selection on other complex traits in humans is inconclusive (86, 87). We suggest that detailed studies of other phenotypes using ancient DNA can be helpful at more generally identifying the types of processes that are important in human evolution.

## Methods

### Capture-shotgun ancient DNA dataset

We downloaded version 37.2 of the publicly available dataset of 2107 ancient and 5637 present-day (Human Origins dataset) samples from the Reich Lab website: https://reich.hms.harvard.edu/downloadable-genotypes-present-day-and-ancient-dna-data-compiled-published-papers. Ancient individuals were treated as pseudo-haploid (i.e. carry only one allele) because most samples are low coverage. For many of these samples, 1,233,013 sites were genotyped using an in-solution capture method (88), while those with whole-genome shotgun sequence data were genotyped at the same set of sites. For our regression models, we included ancient individuals from West Eurasia, defined here as the region west of the Ural Mountains (longitude < 107° E) and north of 35° N (Figure S1). We excluded samples with less than 0.1X coverage. We removed duplicate and closely related samples (e.g. first-degree relatives), by calculating pairwise IBS between ancient individuals. For each individual we identified a corresponding individual with the highest IBS. Based on the distribution of highest IBS values, we identified the pairs with IBS Z-score > 7 as duplicates and retained the individual with fewest missing sites. We also manually removed related samples that were explicitly annotated, selecting the highest coverage representative. Finally, we incorporated additional ancient samples from the Iberian Peninsula from two recent papers (43, 51), applying the same criteria for coverage and relatedness. Ultimately, we compiled a list of 1158 ancient pseudo-haploid individuals in our analyses covering a timespan of the last 40,000 years. Only 11 individuals were dated to >15,000 years BP, and 1147 individuals lived during the last 15,000 years.

### Shotgun ancient DNA dataset

We organized a dataset of samples that were shotgun sequenced without enrichment using the capture array (23, 25–32, 35, 39, 42, 45, 47–51, 54, 56, 58, 60–67). To generate the first 10 PCs from sites on the 1240K capture array, we combined shotgun samples with available genotype data from v37.2 of the Reich lab dataset and 15 individuals that we ourselves pulled down from BAM files. We manually removed duplicate samples in the shotgun dataset, preserving the sample with higher coverage. We checked for first degree relatives in the dataset by considering familial annotations but found none. The final shotgun dataset included 249 individuals.

### Skin pigmentation SNP curation

We obtained summary statistics for UK Biobank GWAS for skin colour (data-field 1717) from the publicly available release by the Neale Lab (version 3, Manifest Release 20180731) (14). The GWAS measured self-reported skin colour as a categorical variable (very fair, fair, light olive, dark olive, brown, black). To identify genome-wide significant and independent SNPs, we performed clumping using PLINK v1.90b6.6 (89) with 1000 Genomes GBR as an LD reference panel (--clump-p1 5 × 10^−8^ --clump-r2 0.05 --clump-kb 250), and followed up with clumping based on physical distance to exclude SNPs within 100 kb of each other. We made two separate lists of UK Biobank SNPs for the shotgun and capture-shotgun datasets because the capture-shotgun dataset was restricted to the 1240K array sites. For the capture-shotgun dataset, we intersected all UK Biobank SNPs with 1240K array SNPs before identifying 170 independent and genome-wide significant SNPs (Table S2), which were used in Figures 1*B*, 1*C*, 2*B*, 2*C*, 3, and 4. For the shotgun dataset, we identified 189 independent and genome-wide significant SNPs (Table S3), which were used in Figures 1*A* and 2*A*.

We also manually curated a list of skin pigmentation-associated SNPs from the literature (Table S4). We identified 12 suitable studies (12, 13, 95, 96, 76, 79, 81, 90–94), 9 of which were GWAS conducted in diverse populations (Table S5). We did not include SNPs from all the considered papers for our analysis because some studies reported associations that were not genome-wide significant (*P* < 5 × 10^−8^). However, we included sub-significant SNPs from a GWAS in Europeans (93) with evidence of replication (*P* < 5 × 10^−8^) in the UK Biobank GWAS. For a GWAS in African populations (12), we selected a SNP for each LD block, which was defined by the study authors, with the greatest support based on multiple lines of evidence (e.g. *P*-value and functional annotations). For a GWAS in South Asians (92), we picked SNPs that were 200 kb away from each other and based the selection of SNPs that were in physical proximity by considering *P*-value. The other studies presented a set of independent SNPs that we considered for inclusion. Lastly, we manually clumped the SNPs curated across studies, selecting SNPs 200 kb apart with more statistical and/or functional evidence than nearby SNPs. Specific details about the choices made on which SNPs to exclude can be found in the “Notes” column for highlighted SNPs in Table S4. We ended with a final list of 18 SNPs for the manually curated set, 10 of which are present on the 1240K capture array (Table S6). We also made pseudo-haploid calls for each of the 18 SNPs in the shotgun dataset, picking a random allele from the reads at each site as for the overall dataset.

### ADMIXTURE analysis on capture-shotgun data

We performed unsupervised ADMIXTURE (97) on the dataset of ancient individuals with K=3, which we found to produce the lowest cross-validation error for 2<K<9. The three identified clusters could easily be identified as corresponding to hunter-gatherer, Early Farmer, and Steppe (also referred to as Yamnaya) ancestry.

### Time series analysis of genetic scores

We calculated genetic scores for each individual from present-day 1000 Genomes populations (64) and ancient individuals. We computed the scores in two ways. First, we weighted by the GWAS-estimated effect sizes. Because of variable coverage across ancient samples, not all SNPs were present in a given sample. To account for missing information in the creation of weighted scores, we devised a weighted proportion in which we divided the realized score over the maximum possible score given the SNPs present in the sample: 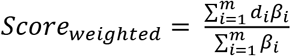 where *m* is the number of skin pigmentation SNPs genotyped in the individual, *d_i_* is the presence of the dark allele at the *i^th^* SNP, and *β*_*i*_ is the GWAS-estimated effect size (Fig. 1 *A-C*). For the manually curated list of SNPs where we did not have comparable effect size estimates, we computed an unweighted score, effectively assuming that all variants had the same effect size. This score is the proportion of dark alleles an individual carries out of the SNPs used in the construction of the score: 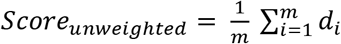 where *m* is the number of SNPs genotyped in the individual and *d_i_* is the presence of the dark allele at the *i^th^* SNP (Fig. 1 *D-F*).

We used logistic regression to examine the association between weighted and unweighted genetic score and time for all ancient samples. We fitted separate models for ancient samples that the shotgun and capture-shotgun datasets. We included ancestry as a covariate in the model by including the first ten principal components. Principal components analysis (PCA) was performed using smartpca v16000 (61) to generate PCs from present day populations and we projected ancient individuals onto these axes of variation. For the shotgun dataset and the capture-shotgun dataset that included all samples (40,000 years BP), we used 1000 Genomes samples. For the capture dataset that restricted to samples later than 15,000 BP, we used West Eurasians from the Human Origins dataset (16).

We model the score of each individual as the proportion of successes in a binomial sample of size *m_i_*. That is, *m*_*i*_*S*_*i*_∼*Binomial*(*m*_*i*_, *p*_*i*_), where, for individual *i*, *m_i_* is the number of SNPs genotyped, *S_i_* is the (weighted or unweighted) score, the probability of success *p_i_* is given by

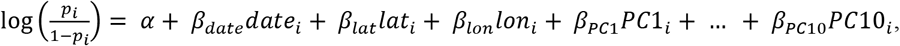

and *date_i_*, *lat_i_* and *lon_i_* are the dates, latitude, and longitude of each individual. For the weighted score, *m*_*i*_*S*_*i*_ is non-integer, but the likelihood can be computed in the same way.

For the stratified analysis, we divided the capture-shotgun dataset into mutually exclusive ancestry groups. We categorized individuals for this analysis as hunter-gatherer if they carried over 60 percent of the ADMIXTURE-estimated hunter-gatherer component. For placement into the Early Farmer group, we required that an individual carry over 60 percent of that component. Individuals we categorized as Steppe had over 30 percent of the ADMIXTURE component and were dated to less than 5000 years BP. Based on these cutoffs, there were 102 hunter-gatherer, 499 Early Farmer, and 478 Steppe ancestry assigned individuals.

Because the fitted model parameters are overdispersed (Fig. S2), we computed *P*-values from a genome-wide empirical null distribution for *β_date_*. We made this null distribution by running the regression model described above on scores from sets of random, frequency-matched (+/− 1%) SNPs across the genome, maintaining the same effect sizes for the weighted score. We matched the derived allele frequency based on the EUR superpopulation frequencies from 1000 Genomes. We reported *P*-values based on 10,000 random samples.

### Time series analysis of individual SNP allele frequencies

We performed logistic regression for each individual SNP using date of the sample and ancestry as covariates. That is, the probability that the haplotype sampled from individual carries the derived allele is *p_i_*, where

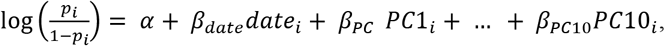

We compared the full model above to a nested model with no principal components by performing a likelihood ratio test (in R we use anova(nested model, full model, test=‘Chisq’)) to obtain a *P*-value for the ancestry term which encompasses *β_PC1_*…*β_PC10_*. To obtain a *P*-value for *β_date_*, we compare a nested model without *β_date_* to the full model.

### Q_x_ polygenic selection test

We used the *Q_x_* test for directional, polygenic selection on skin pigmentation (10). For this test, we used skin pigmentation SNPs obtained from the UK Biobank in all 1000 Genomes populations. We restricted sites to those on the 1240K and constructed a covariance matrix from a total of 1 million SNPs. To calculate empirical *P*-values, we sampled a total of 500,000 null genetic values matching skin pigmentation SNPs based on a +/− 2% frequency on the minor allele. However, we report 1 million runs for the top 7 SNPs and 2 million runs for the top 5 and 6 to better distinguish the deviation of the tested SNP sets from the expectation under drift.

### Variance in skin pigmentation explained by individual SNPs

We estimated the variance explained in the phenotype by a SNP within the UK population using

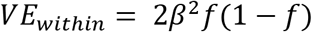

where *β* is the UK Biobank GWAS-estimated effect size and *f* is the allele frequency in the GBR population from 1000 Genomes. We calculated the proportion of variance explained as

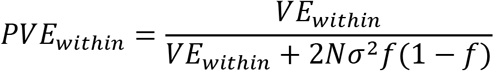

where *N* is the GWAS sample size (356,530) and σ is the standard error of the estimated *β* (98). To estimate the variance explained between GBR and YRI, we calculated the between population variance as

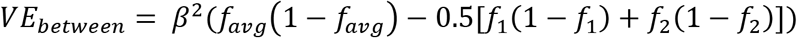

where *f_1_* and *f_2_* are the allele frequencies of GBR and YRI, *β* is the UK Biobank GWAS-estimated effect size, and *f_avg_* is 0.5(*f_1_* + *f_2_*).

### Population branch statistic

We calculated the population branch statistic (PBS) (63) for different sets of ancient groups and present-day groups. To obtain groups with ancient individuals corresponding to the three ancestral populations of Europeans, we used the ADMIXTURE inferred ancestry proportions. Individuals were included in the hunter-gatherer and Early Farmer groups based on having ≥80% global ancestry. For Yamnaya or Steppe, we required ≥ 60% ancestry and an estimated date <6500 years BP. Based on these cutoffs (more stringent than the ones used for the stratified time series analysis), there were 75 hunter-gatherer, 241 Early Farmer, and 81 Steppe ancestry assigned individuals.

Using the allele frequencies from these ancient groups and present-day 1000 Genomes populations, we estimated pairwise Hudson’s F_ST_ between groups for the PBS test. The estimator is calculated as

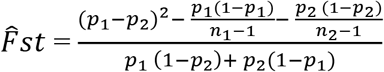

where for population *i*, *n_i_* is the sample size and *p_i_* is the allele frequency of the sample. SNPs were excluded if a sample group did not contain at least 10 individuals. Because of variable coverage in ancient genomes, not all sites were used in the calculation. *F_ST_* was calculated using 20 and 40-SNP non-overlapping windows throughout the genome. *F_ST_* values were transformed to calculate genetic divergence between populations as *T* = −log(1 − *F_ST_*). We calculated PBS for ancient group *X* (i.e. hunter-gatherers, Early Farmers, or Steppe) using 1000 Genomes Han Chinese (CHB) and Yoruba (YRI) as

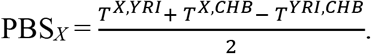

We identified nearby genes by examining the NCBI RefSeq track for assembly GRCh37/hg19 (downloaded from https://genome.ucsc.edu/cgi-bin/hgTables) for genes that overlap with windows with high PBS values for hunter-gatherers or Early Farmers.

## Code availability

Scripts used to generate the main results and figures of this paper are available at https://github.com/mathilab/SkinPigmentationCode.

## Acknowledgments

We thank Sarah Tishkoff, Alexander Platt, Bárbara Bitarello, Samantha Cox, Arslan Zaidi, and Andrew Chen for helpful comments on earlier versions of this manuscript. This research was supported by a Research Fellowship from the Alfred P. Sloan foundation [FG-2018-10647], a New Investigator Research Grant from the Charles E. Kaufman Foundation [KA2018-98559], NIGMS award number [R35GM133708], and NRSA T32 [T32GM008216]. The content is solely the responsibility of the authors and does not necessarily represent the official views of the National Institutes of Health.

