## Supplemental figures and supplemental table 1 for "The evolution of skin pigmentation associated variation in West Eurasia"

### Supplementary Figures

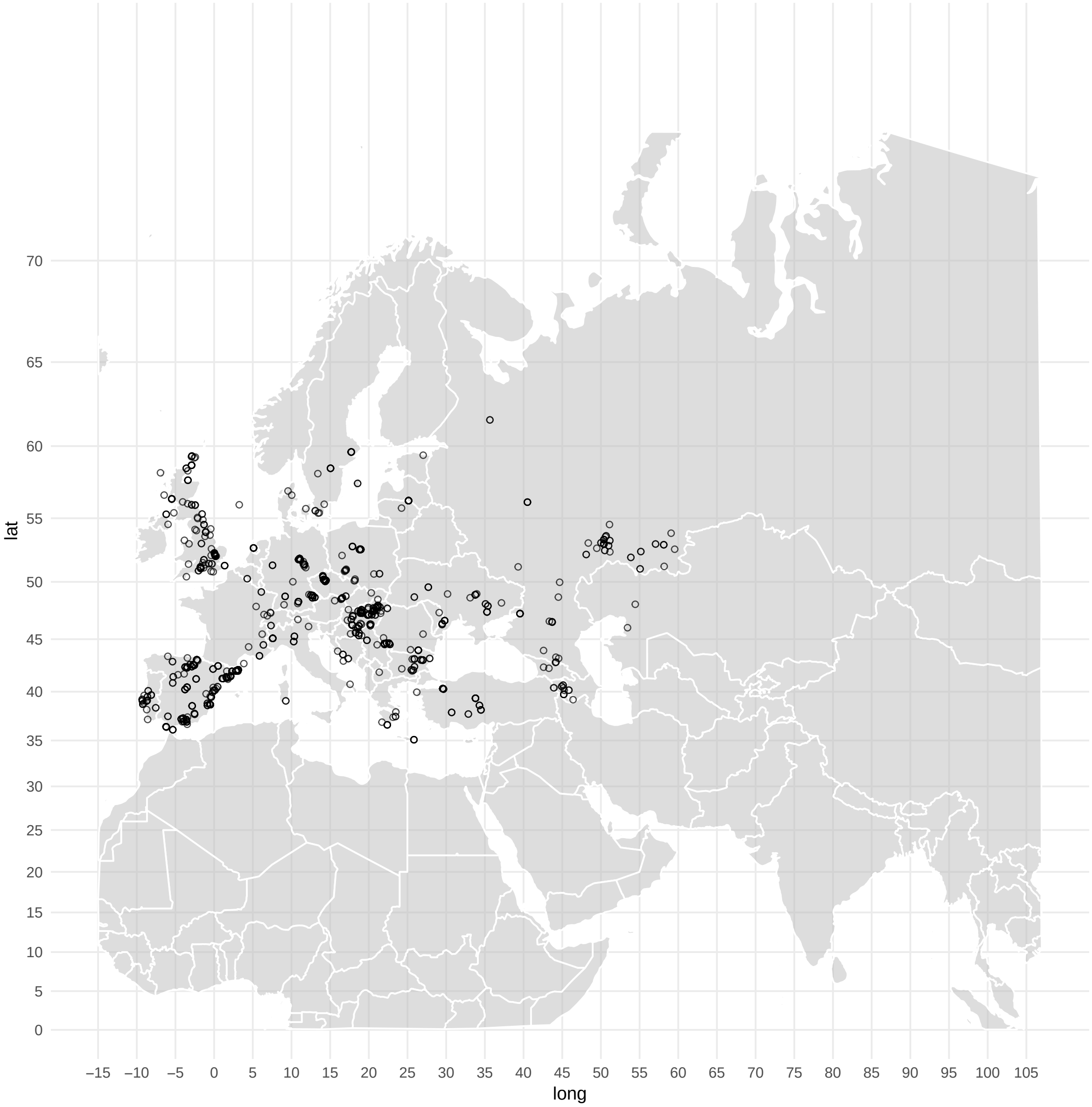


**Supplementary Figure S1**. Map of locations of ancient individuals included in the final capture-shotgun dataset.

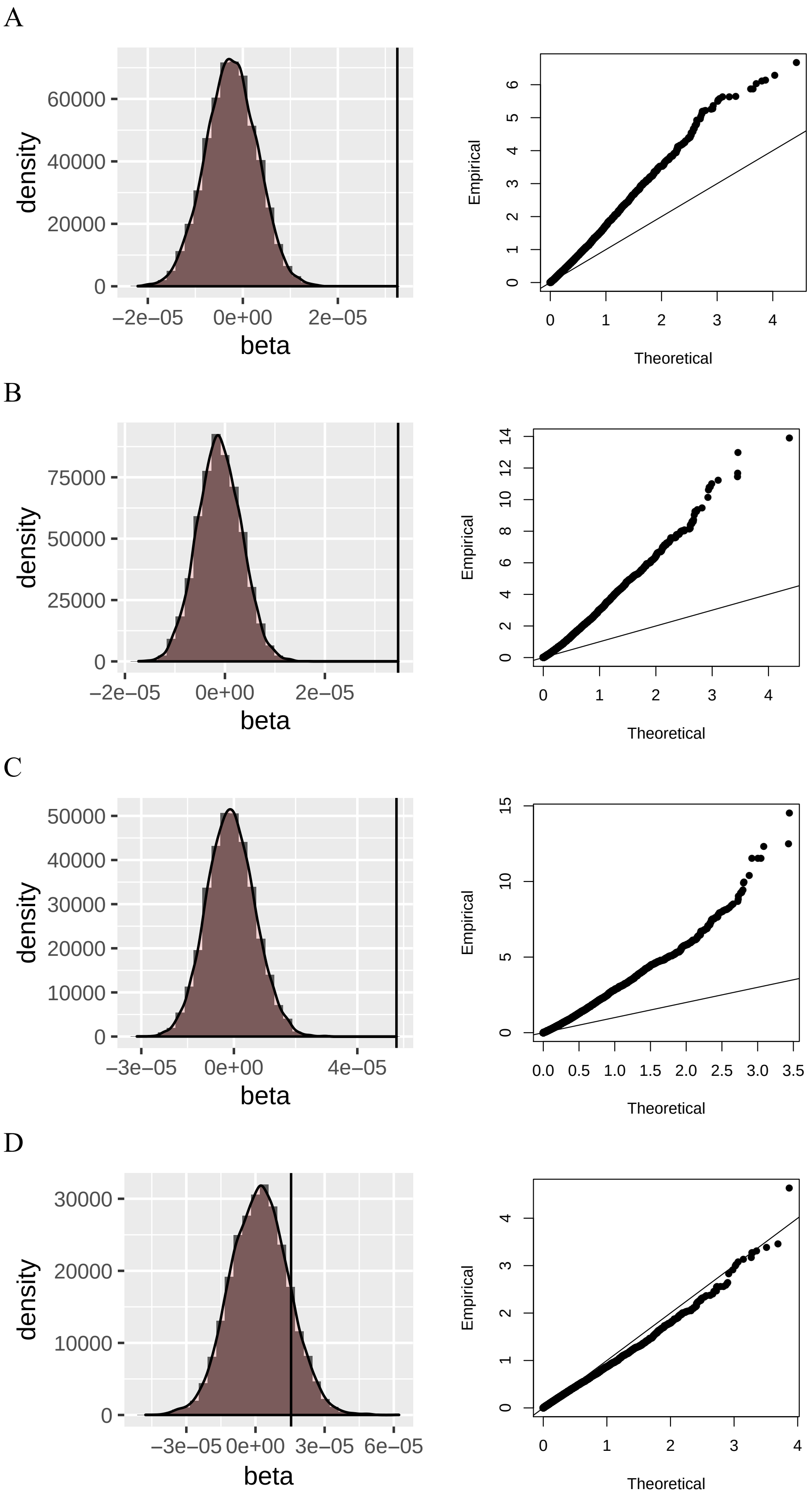
A
B
C
D
density
beta
Empirical
Theoretical
60000
40000
20000
0
−2e−05
0e+00
2e−05
75000
50000
25000
50000
40000
30000
20000
10000
−3e−05
4e−05
30000
3e−05
6e−05

E

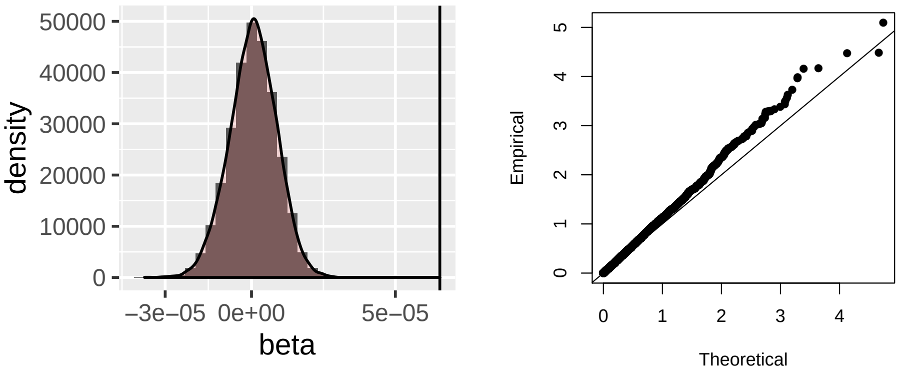


F

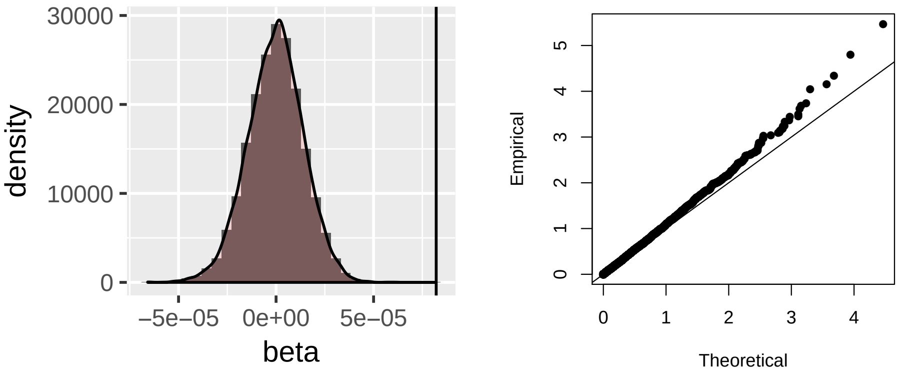


**Supplementary Figure S2**. Distribution of $\beta_{date}$ from regression model for randomly generated score over time regressions and Q-Q plot of -log$_{10}$(P-value) of $\beta_{date}$. Randomly generated scores based on UK Biobank SNPs and manually curated SNPs were calculated for samples in the (A,D) shotgun dataset, (B,E) capture-shotgun dataset, and (C,F) capture-shotgun dataset dated within the past 15,000 years.

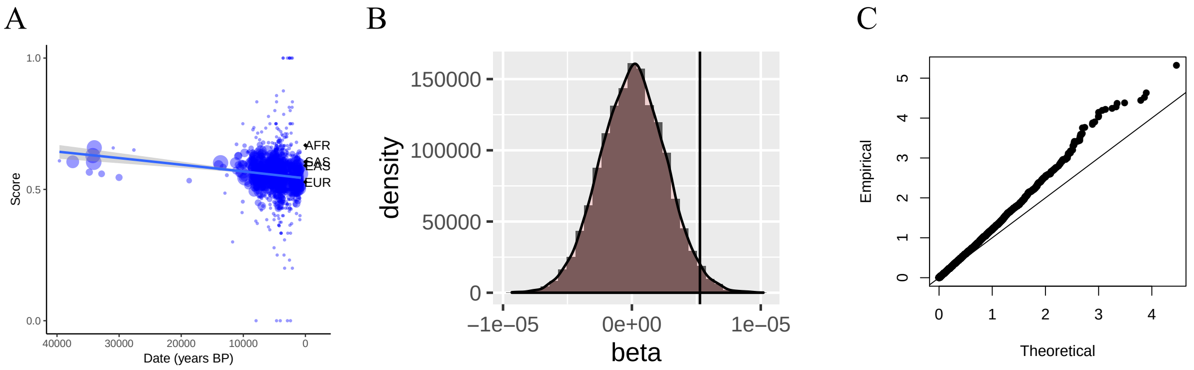


**Supplementary Figure S3**. (A) Regression of unweighted score based on 170 UK Biobank SNPs over time. (B) Distribution of $\beta_{date}$ from regression model for randomly generated score over time regressions and (C) corresponding Q-Q plot of -$\log_{10}$($P$-value).

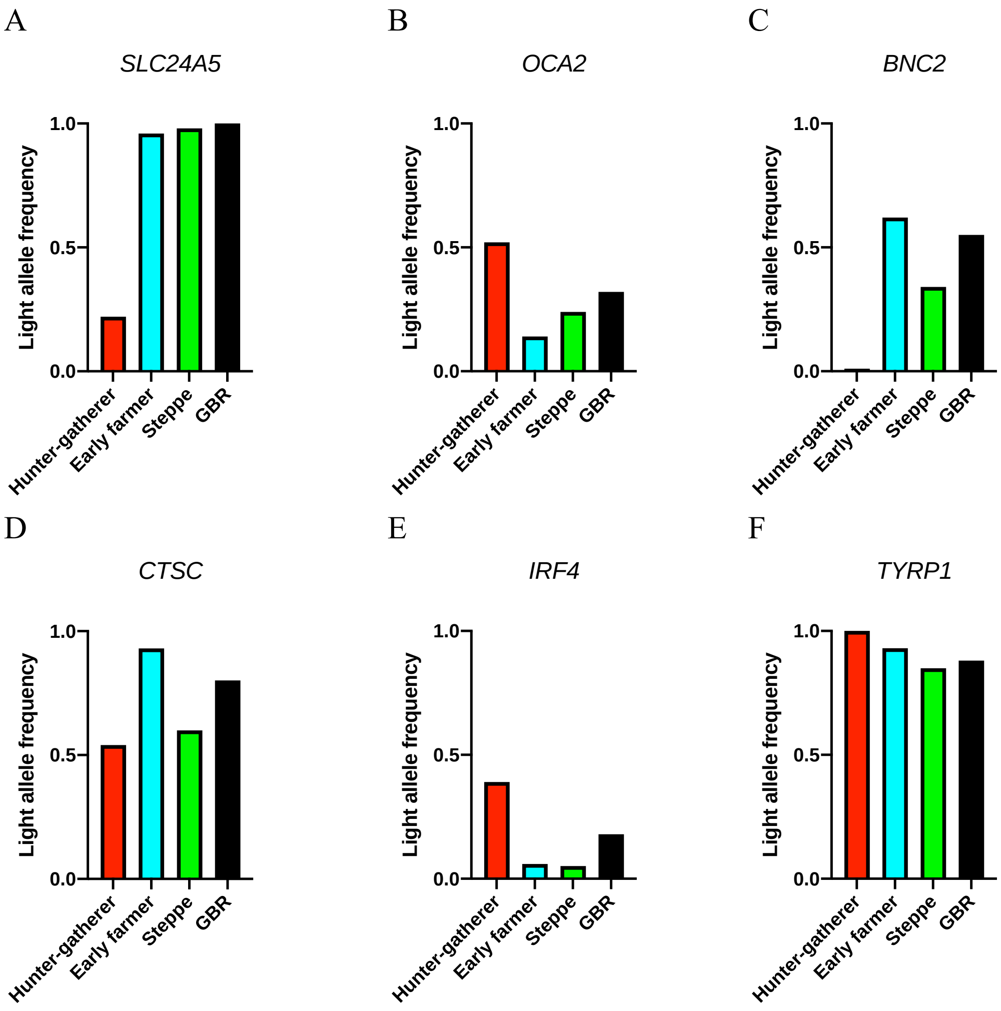


**Supplementary Figure S4**. Light allele frequencies in ancient and present-day European populations for select SNPs (A) rs2675345, (B) rs4778123, (C) rs2153271, (D) rs3758833, (E) rs12203592, and (F) rs1325132.

A

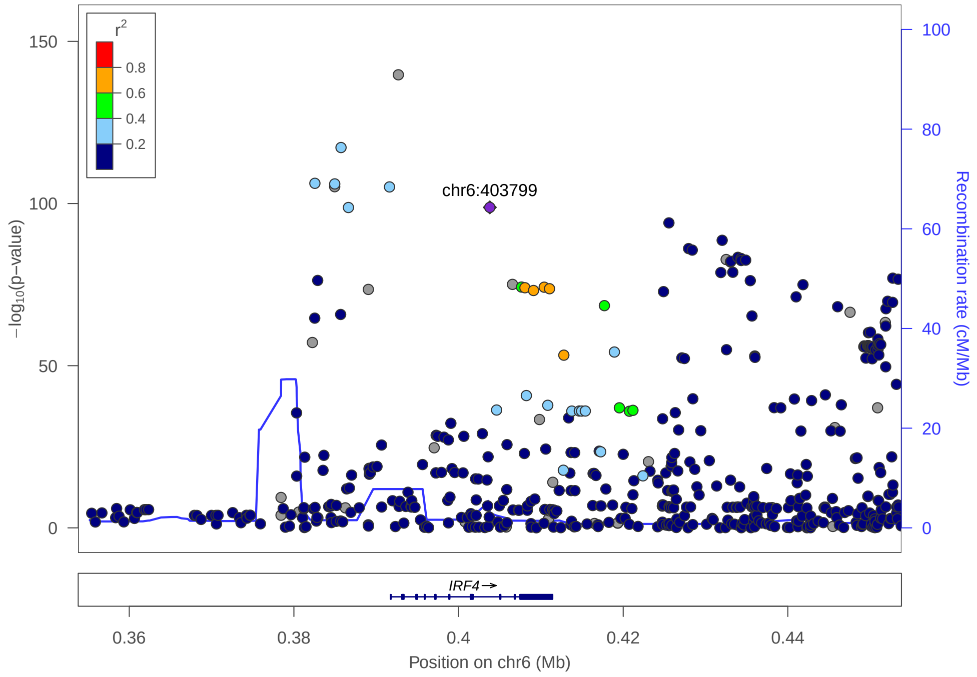


B

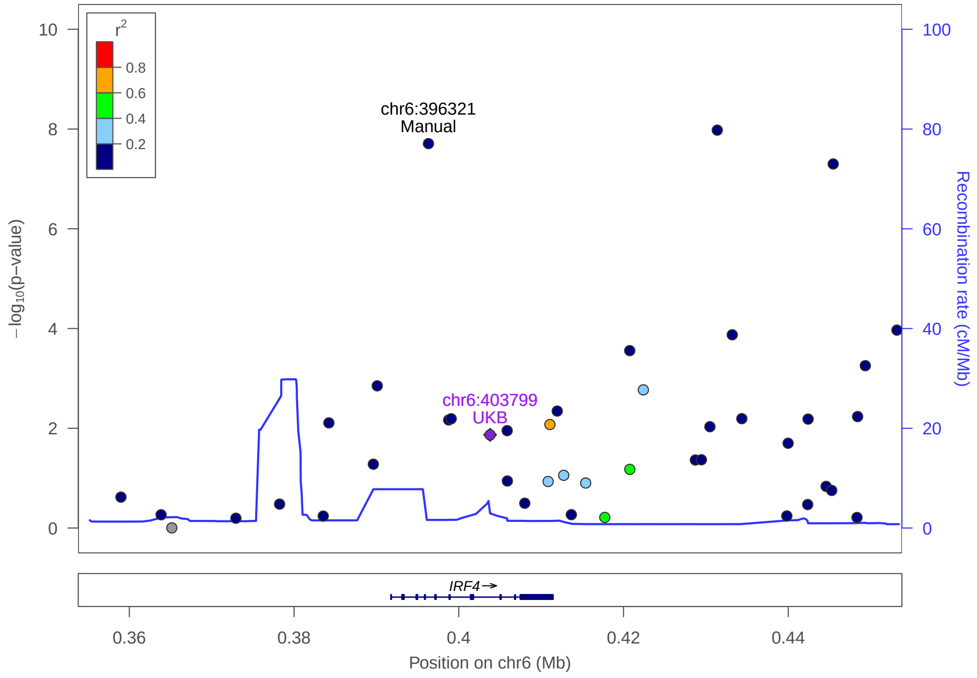


**Supplementary Figure S5.** (A) LocusZoom plot of the *IRF4* region for UK Biobank SNPs with the y-axis reporting P-values for the SNP association with skin colour association. Note: Low-confidence variants are included, but for SNP selection low-confidence variants were removed. (B) Plot of 1240K array SNPs at the *IRF4* locus with P-values corresponding to the ancestry term in the regression model for capture-shotgun 40,000 years with the manually curated SNP.

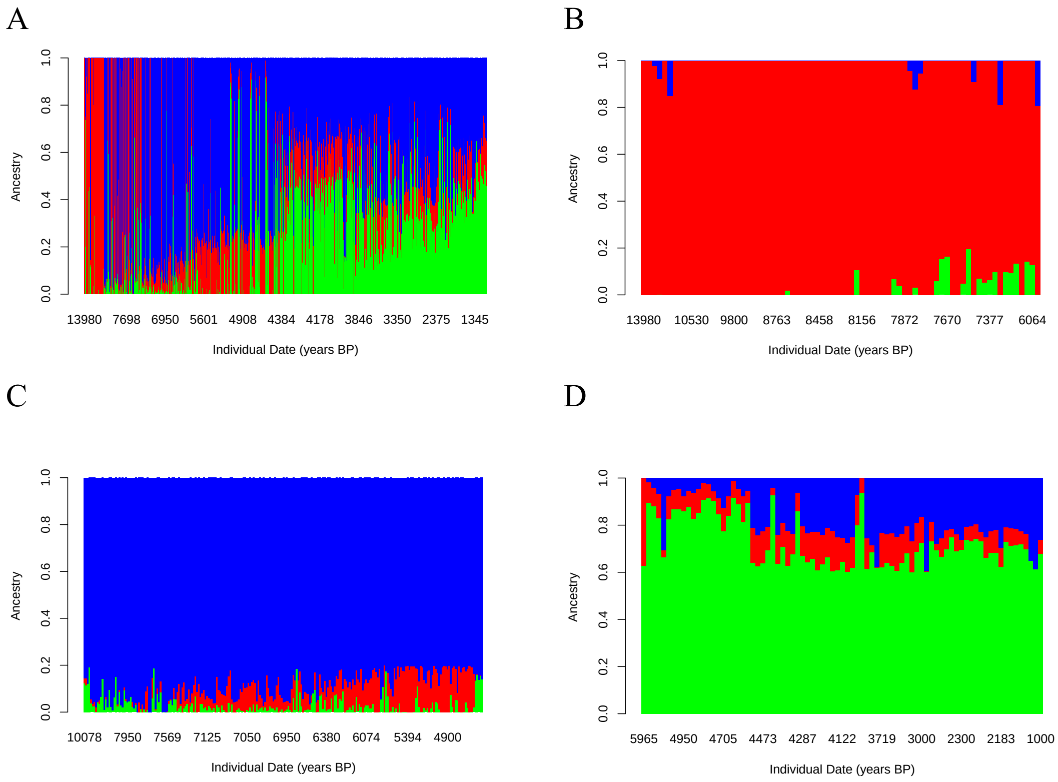


**Supplementary Figure S6**. Unsupervised admixture results (K=3) for (A) all capture-shotgun ancient samples from 15,000 years BP onward, samples categorized for PBS analysis as (B) Hunter-gatherer (red component ≥ 0.8), (C) as Early Farmer (blue component ≥ 0.8), and (D) as Steppe (green component ≥ 0.6 and estimated date < 6500 years BP). The samples for each population were used in the PBS analysis.

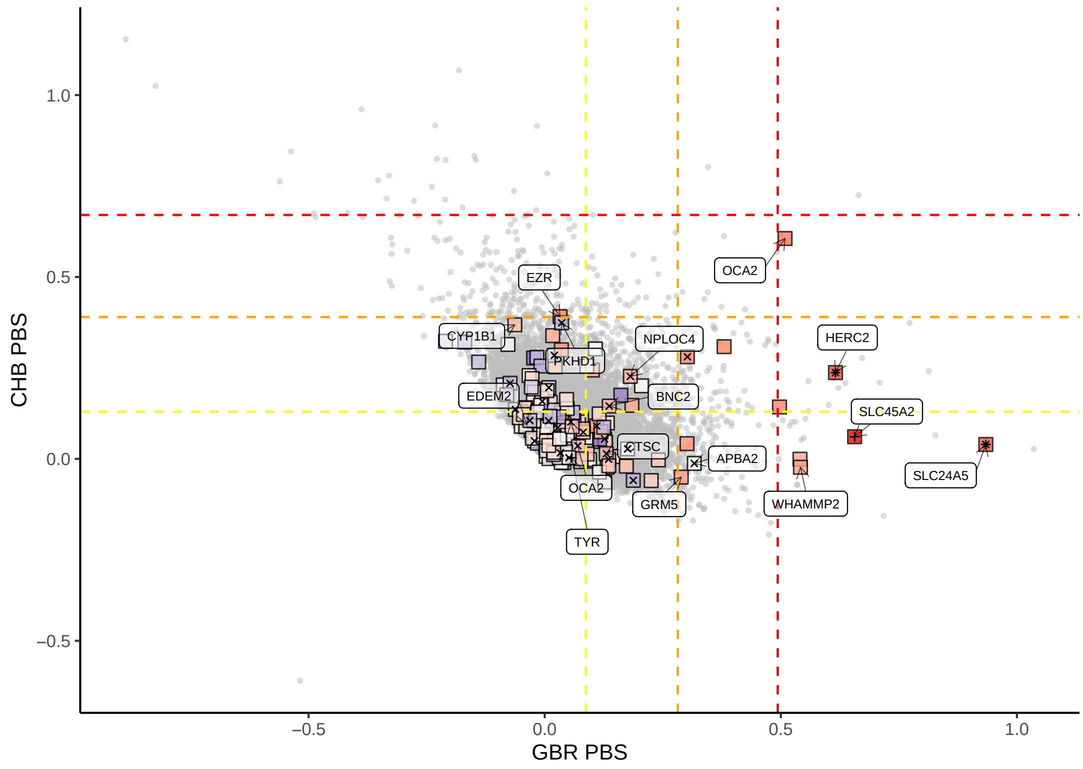


**Supplementary Figure S7**. Joint distribution of PBS for CHB and GBR from 1000 Genomes for the tree of GBR-CHB-YRI.

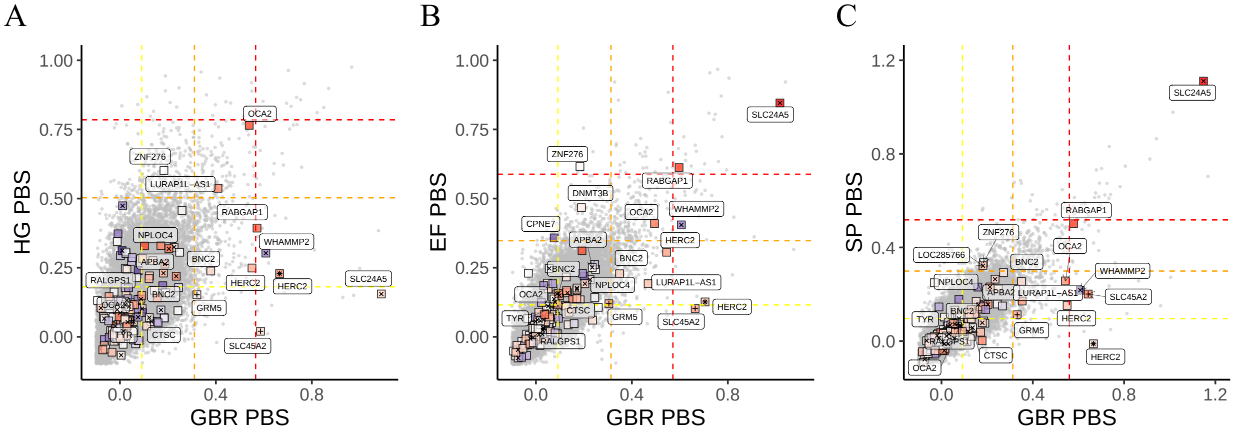


**Supplementary Figure S8**. Joint PBS distributions for non-overlapping 20 SNP-windows across the genome for GBR and (A) Hunter-gatherer, (B) Early Farmer, (C) Steppe using YRI and CHB. Boxes represent windows centered around a UK Biobank skin pigmentation SNP and are colored according to the magnitude of difference in frequency between the ancient group and YRI. Red indicates the light allele is higher in frequency in the ancient group. Plusses (+) indicate whether a window contains a skin pigmentation SNP at the extremes of the time term distribution from Figure 2, while crosses (x) indicate a variant at the extreme of the ancestry term distribution. Nearest genes are labelled. Yellow, orange, and red lines represent top 20, 1, and 0.1 percentiles.

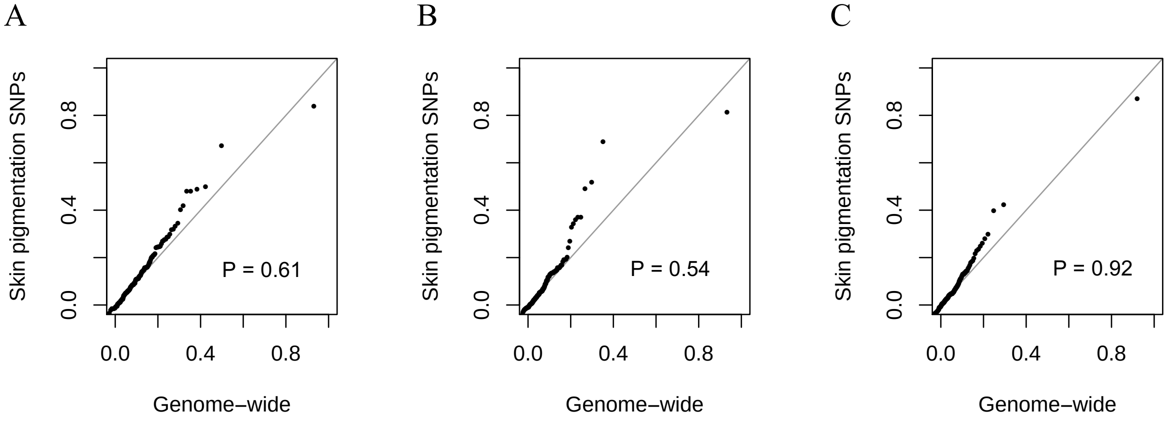


**Supplementary Figure S9**. Q-Q plots of distributions of PBS at skin pigmentation SNP-harboring genomic windows of 40 SNPs and PBS of all genomic windows for (A) hunter-gatherers, (B) early farmers, and (C) Steppe.

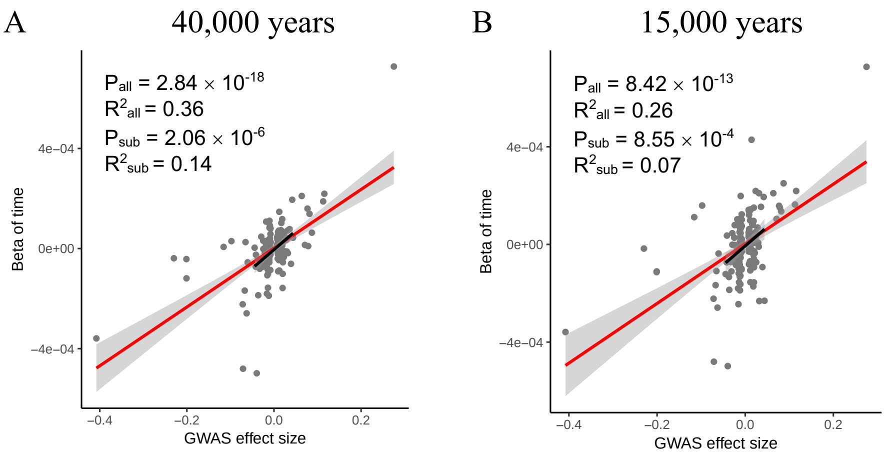


**Supplementary Figure S10**. Plots of 170 UK Biobank SNP GWAS effect size and magnitude of change in allele frequency represented by $\beta_{date}$ in the full regression model for capture dataset over (A) 40,000 and (B) 15,000 years. Regression lines in red are based on 170 SNPs and statistics for this regression are denoted by "all," whereas the black line is based on 147 smaller GWAS effect size SNPs ($|\beta_{date}| < 0.05$) and statistics denoted by "sub."

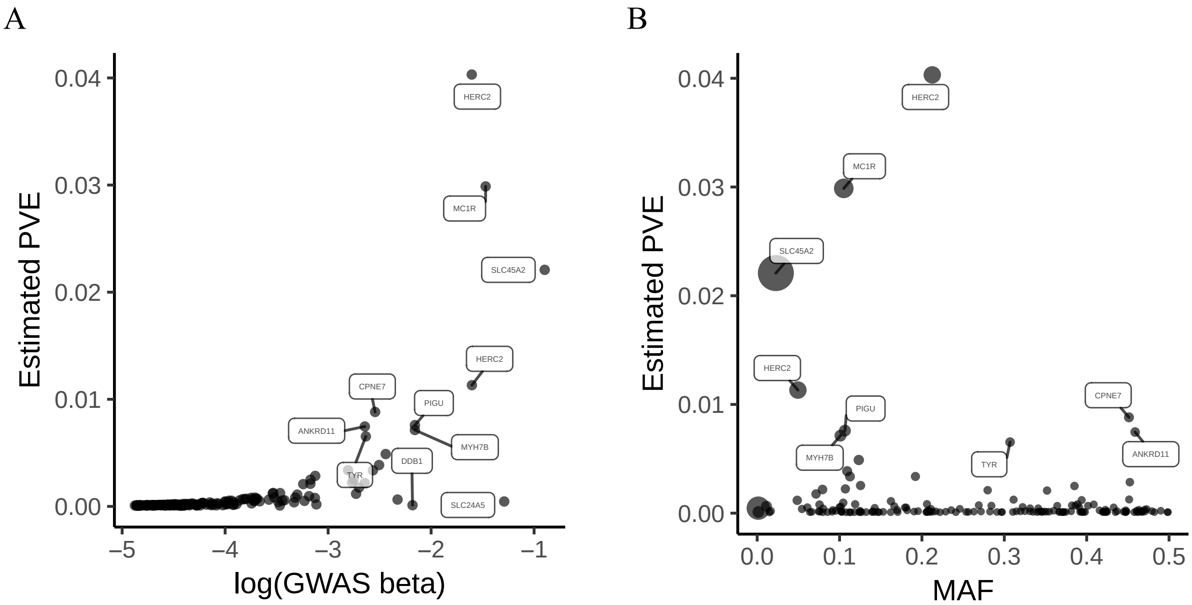


**Supplementary Figure S11**. (A) Estimated proportion of variance explained in the UK Biobank European population over GWAS estimated effect size of the 170 skin pigmentation SNPs. (B) Estimated proportion of variance explained over minor allele frequency in the UK Biobank European population. Point areas are scaled by effect size.

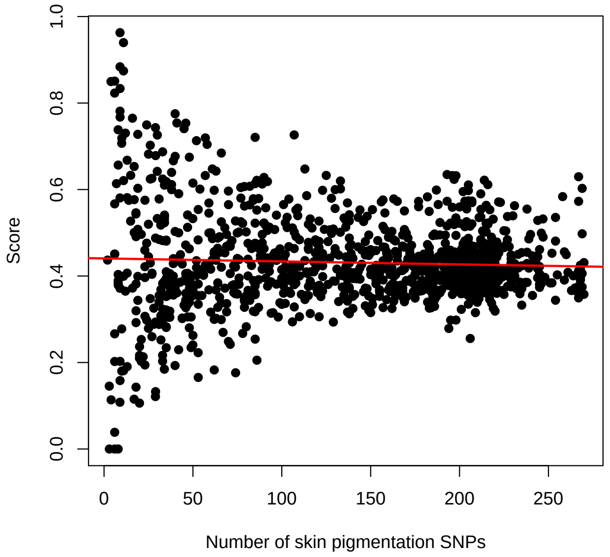


**Supplementary Figure S12**. Regression of skin pigmentation genetic score regressed on number of UK Biobank skin color SNPs present.

| Name | UK Biobank SNP score | UK Biobank SNPs present | Manually curated SNP score | Manually curated SNPs present |
|---|---|---|---|---|
| Altai | 0.81 | 169 | 0.67 | 18 |
| Denisova | 0.80 | 140 | 0.61 | 18 |
| Les Cottés | 0.86 | 59 | 0.71 | 17 |
| Goyet | 0.75 | 81 | 0.69 | 16 |
| Mezmaiskaya 1 | 0.78 | 66 | 0.75 | 4 |
| Mezmaiskaya 2 | 0.91 | 62 | 0.75 | 16 |
| Spy | 0.74 | 93 | 0.57 | 7 |
| Vindija | 0.84 | 153 | 0.67 | 18 |

**Supplementary Table S1**. Weighted and unweighted genetic scores based on 170 UK Biobank SNPs and 18 manually curated SNPs, respectively, for Denisovan and Neanderthals.
